# The Demographic History of Micro-endemics: Have Rare Species Always Been Rare?

**DOI:** 10.1101/2020.03.10.985853

**Authors:** Andrew J. Helmstetter, Stuart Cable, Franck Rakotonasolo, Romer Rabarijaona, Mijoro Rakotoarinivo, Wolf L. Eiserhardt, William J. Baker, Alexander S.T. Papadopulos

## Abstract

Extinction has increased as human activities impact ecosystems. Conservation assessments for the IUCN red list are a fundamental tool in aiding the prevention of further extinction, yet, relatively few species have been thoroughly assessed. To increase the efficiency of assessments, novel approaches are needed to highlight threatened species that are currently data deficient. Many Madagascan plant species currently have extremely narrow ranges, but this may not have always been the case. To assess this, we used high-throughput DNA sequencing for 2-5 individuals of each species - reflecting the paucity of samples available for rare species. We estimated effective population size (*N*_*e*_) for each species and compared this to census population *(N*_*c*_) sizes when known. In each case, *N*_*e*_ was an order of magnitude larger than *N*_*c*_ – a signature of rapid, recent population decline. We then estimated the demographic history of each species, tracking changes in *N*_*e*_ over time. Five out of ten species displayed significant population declines towards the present (68–90% decreases). Our results for palm trees indicate that it is possible to predict extinction risk, particularly in the most threatened species. We performed simulations to show that our approach has the power to detect population decline during the Anthropocene, but performs less well when less data is used. Similar declines to those in palms were observed in data deficient species or those assessed as of least concern. These analyses reveal that Madagascar’s narrow endemics were not always rare, having experienced rapid decline in their recent history. Our approach offers the opportunity to target species in need of conservation assessment with little prior information, particularly in regions where human modification of the environment has been rapid.

**Summary:** Current IUCN conservation assessment methods are reliant on observed declines in species population and range sizes over the last one hundred years, but for the majority of species this information is not available. We used a population genetic approach to reveal historical demographic decline in the rare endemic flora of Madagascar. These results show that it is possible to predict extinction risk from demographic patterns inferred from genetic data and that destructive human influence is likely to have resulted in the very high frequency of narrow endemics present on the island. Our approach will act as an important tool for rapidly assessing the threatened status of poorly known species in need of further study and conservation, particularly for tropical flora and fauna.

## Introduction

The existence of a mass extinction during modern times has become well accepted (Ceballos *et al*. 2017). According to the International Union for Conservation of Nature (IUCN) red list, approximately 1.2% of assessed species are extinct in the wild and 29.7% are threatened by extinction (Pimm *et al*. 2015). With fewer than 100k species assessed, we have poor knowledge of the extinction risk in many taxa, including those that are vital to human wellbeing. For example, only 27k plant species have IUCN assessments (∼6% of all plants), but the loss of plant diversity is expected to have a greater effect on humans than the loss of diversity in any other group (Schatz 2009). Therefore, it is critical that we assess and continue to monitor plant species. Despite the rigour with which assessments are made, it is possible that current methods substantially underestimate the extent to which species may be at risk of extinction. For example, population decline within the last 100 years is a fundamental component of how IUCN red listing is performed (Maes *et al*. 2015; Collen *et al*. 2016), but recent estimates suggest that 30% of all vertebrates species are in decline and as much as 55% of declining bird species are not classified as threatened (Ceballos *et al*. 2017). Furthermore, substantial population decline in many species may have pre-dated historical records or may not have been observed directly at all. As a result, it is unclear whether species that are rare today have experienced population decline or simply have a low carrying capacity (e.g. after becoming highly specialised on spatially-restricted habitats). Without taking into account unobserved declines in population sizes we may be drastically underestimating the number of threatened species and the severity of the extinction threat they face. The unique and highly endemic flora of Madagascar provide an opportunity to investigate the link between historical demographic change and extinction risk.

Levels of endemism on Madagascar are some of the highest worldwide; all of the island’s amphibians and terrestrial mammals, 83% of vascular plant species and 86% of macroinvertebrates are found nowhere else (Allnutt *et al*. 2008; Vences *et al*. 2009; Goyder *et al*. 2017). A striking feature of the Madagascan flora and fauna is the abundance of micro-endemic species that have small ranges and are relatively rare (Yoder *et al*. 2000; Wilme *et al*. 2006; Vences *et al*. 2009). For example, more than half of the island’s endemic palms are known to be extremely rare – many are found in just a single population or with fewer than 100 individuals in the wild (Rakotoarinivo *et al*. 2014). The processes that gave rise to the incredible diversity and micro-endemism of Madagascar are hotly debated (Wilme *et al*. 2006; Pearson & Raxworthy 2009; Vences *et al*. 2009; Eberhart-Phillips *et al*. 2015; Federman *et al*. 2015; Yoder *et al*. 2016; Hackel *et al*. 2018), but proposed models put little emphasis on the role human beings have played in shaping the distribution of species. Humans arrived on Madagascar recently, around 2500 years ago, and preceded the rapid extinction of the island’s megafauna (Crowley 2010). Although the exact causes of megafaunal extinction are less clear, increasing human activities (including hunting, slash and burn agriculture, illegal logging and introduction of invasive species) are likely to have played a major role (Godfrey *et al*. 2019). Extinction of large lemuriforms is also thought to have had a negative impact on dispersal and fitness of large-seeded plants (Federman *et al*. 2016). Deforestation has also advanced rapidly so that only 10% of the island’s forest remains (Harper *et al*. 2019). This combination of recent human influence and abundant micro-endemism invites the question: have Madagascar’s micro-endemics always been rare, or is this pattern driven by exceptional population decline across the island’s flora and fauna? In this study, our aims were to (i) investigate whether Madagascan plants have suffered population declines in the recent past and (ii) determine whether inference of past demographic changes from genetic data can predict extinction risk.

## Results and Discussion

We investigated population decline in 10 Madagascan plant species using genome-wide genetic markers. Our sample included a range of endemic species from the humid and littoral forests of eastern Madagascar (Fig. 1). These forests have decreased by up to a third since the 1970s and are severely under threat (Moat & Smith 2007). The diverse palm (Arecaceae) flora of Madagascar is reasonably well studied and more than 90% of the 204 Madagascan palms have been placed in one of the threatened categories following red list assessment – an indication that these all have declining or small populations. For this reason, we included four palm species (all of which have decreasing populations) as our training data set. Two of these have been assessed as near threatened -*Ravenea robustior* (Rakotoarinivo & Dransfield 2012b) is estimated to have <1000 mature individuals in the wild, while *Dypsis procumbens* is more common but restricted to 32 locations and is exploited for timber (Rakotoarinivo & Dransfield 2012a). *Dypsis rabepierrei* is critically endangered with no more than 20 individuals known from a single location (Eiserhardt *et al*. 2018) and *Satranala decussilvae* is endangered with fewer than 200 mature trees in the wild (Rakotoarinivo & Dransfield 2012c).

**Figure 1.**
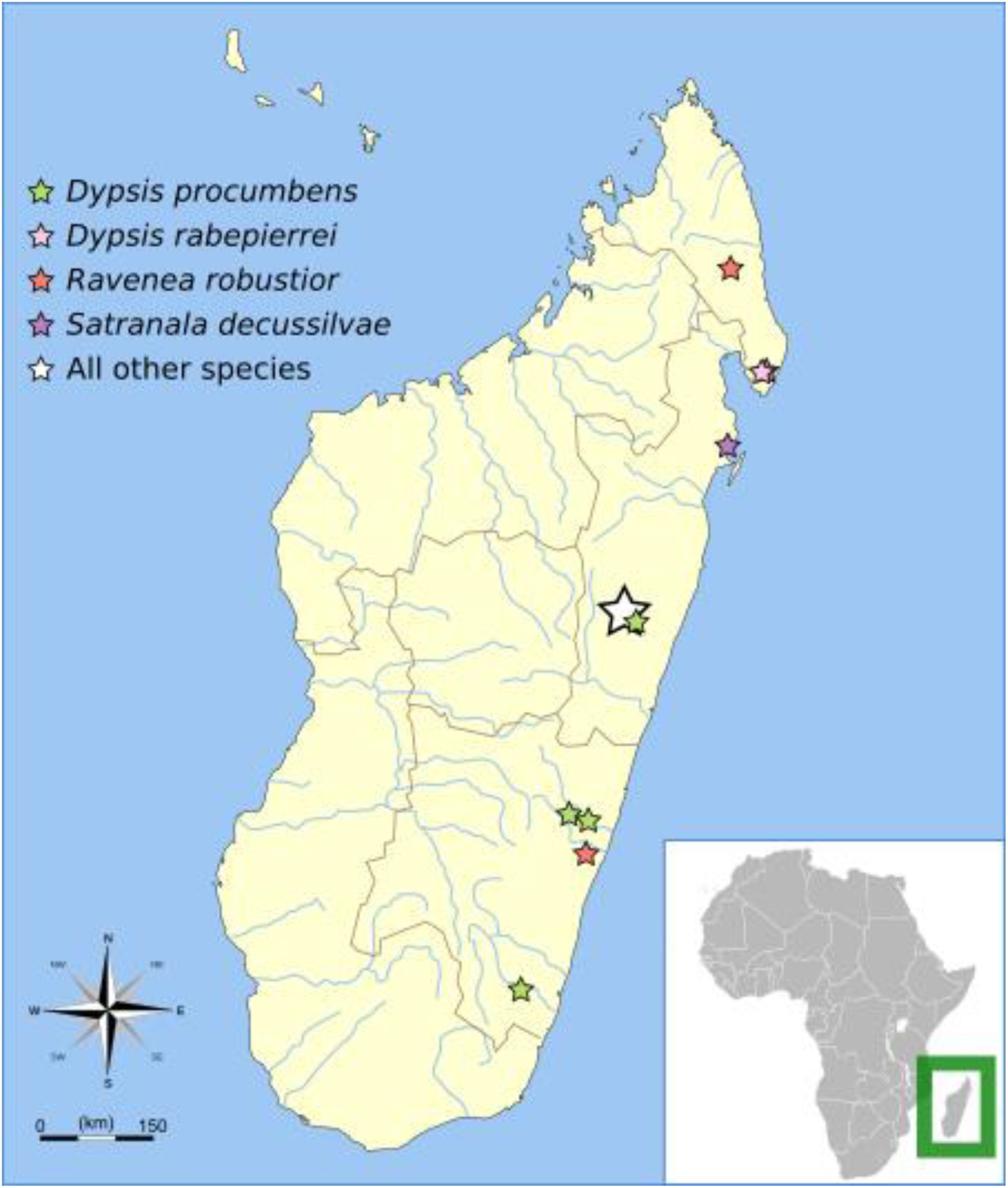
A map of collection sites for the 10 species used in this study. Stars represent approximate collection locations and are colour coded for each palm species sampled. The remaining six species were collected from a single location shown as a white star. Original map of Madagascar by Maky and shared under the CC BY-SA 3.0 license (https://creativecommons.org/licenses/by-sa/3.0/deed)

The effective population size (*N*_*e*_) of a species is normally substantially smaller than the census population size, such as in humans where *N*_*e*_ is in the order of 10,000 (Gazave *et al*. 2013). When populations are experiencing rapid decline it is possible for *N*_*e*_ to substantially exceed census population size, as has been observed in the critically endangered black rhinoceros (Braude & Templeton 2009). If the studied palm species have always been rare, we expected that *N*_*e*_ would be equal or less than the census population sizes. Conversely, if rare species were more abundant prior to human influences in Madagascar *N*_*e*_ would be substantially higher than contemporary census sizes. We performed double digest restriction site associated DNA (ddRAD) sequencing (Peterson *et al*. 2012) on 2-5 individuals of each species and genotyped these at 24-78k loci using the Stacks (Catchen *et al*. 2013) pipeline (see table S1 and Supplementary materials and methods for details). We then estimated *θ*, the population scaled mutation rate (*θ* = 4*N*_*e*_*µ*), using the Bayesian method implemented in the R package *thetamateR*, which allows for rate variation between loci. The estimated *θ* value for each species was converted to *N*_*e*_ assuming the average mutation rate in angiosperms (De La Torre *et al*. 2017) and a generation time of 10 years. In each the four palm species, estimated *N*_*e*_ substantially exceeds the current census population sizes (Fig. 2, table S1), a strong indication that these rare species have not always been rare.

**Figure 2.**
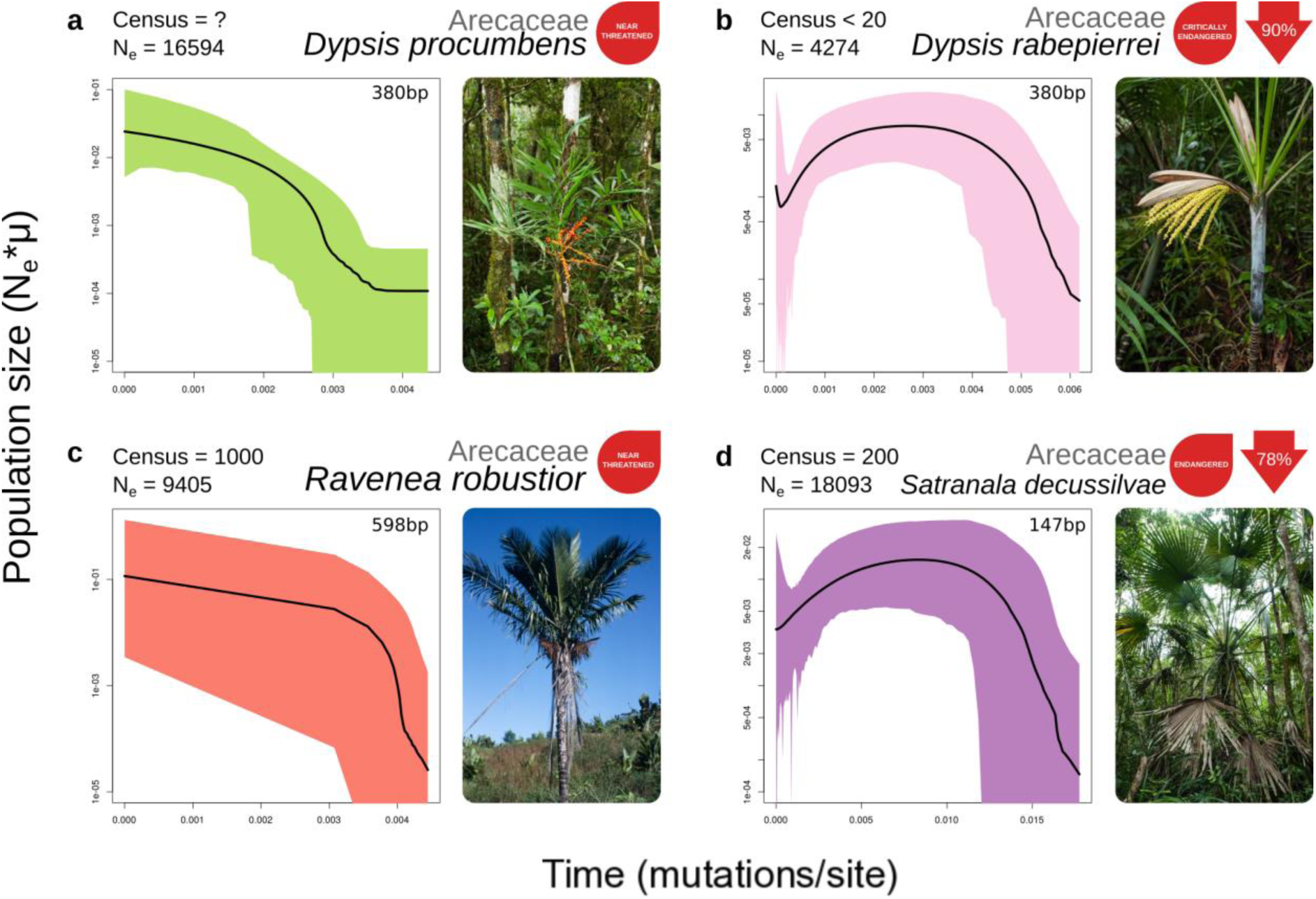
Effective population size through time for four species of palm that have conservation assessments. The present day is on the left side of each plot. Black lines represent the median population size and shaded polygons show the 95% highest posterior density (HPD). To the right of each plot is a picture of the study species and the current IUCN red list assessment. Values in red arrows indicate the percentage decline from maximum population size to the minimum population size inferred after the maximum. The increase in the median value of population size for *D. rabepierrei* in the very recent past is the result of a lack of coalescent events, which is reflected by extremely wide 95% HPD values during this period, and should not be interpreted as population size increase. Results of skyline plots for simulated data are shown as coloured Photo of *R. robustior* by Henk Beentje. All other photos by WJB.

To provide further evidence that these palm populations have experienced declines, we used a subset of loci to investigate the demographic history through time using the Extended Bayesian Skyline Plot approach (EBSP; Heled & Drummond 2008) implemented in BEAST v2.4.2 (Bouckaert *et al*. 2014) following the methodology outlined by Trucchi et al. (Trucchi *et al*. 2014). These methods struggle to estimate *N*_*e*_ in the very recent past due to a paucity of coalescent events and maintenance of variation in diminished populations, so declines very close the present may not be inferred. Given that our *N*_*e*_ estimates and IUCN assessment suggest our four palm species have rapidly decreasing populations, we expected that species population sizes over time (estimated using EBSP) would decrease sharply towards the present. The endangered palm species have very small census sizes, so we expected that any decline would be more pronounced in these species. If so, this approach could provide a useful indicator of extinction risk, the efficacy of which can be demonstrated using our palm dataset.

For each species, we assessed whether we could reject a hypothesis of constant size through time by assessing the “sum(indicators.alltrees)” statistic, which is analogous to the number of population size change events. When 95% highest posterior density (HPD) values do not include zero this indicates that *N*_*e*_ has not been constant through time. The results of the EBSP analysis for palms (Fig. 2) do not contain evidence of a recent decline in the near threatened species, each with a single population size change event, corresponding to early population expansion (Table S1). It is possible that these species at less risk are more abundant than thought. However, given the large difference between *N*_*e*_ and census sizes (Table S1), this could also indicate a very rapid a steep decline in recent history that could not be detected with EBSP. On the other hand, the endangered species have experienced very steep population declines towards the present, as much as a 90% reduction in population size, and multiple demographic events in their history. These results are expected for an endangered species, so we suggest that the approach used here is a useful tool for predicting more extreme levels of extinction risk.

To validate whether our approach has the power to detect anthropogenic population decline, we simulated sequence datasets using *fastsimcoal2* (Excoffier et al. 2013) with ancestral populations sizes set to *thetamateR* Ne estimates and a range of population declines of varying severity that took place 2,500 years ago. We then ran our pipeline, sampling loci for the simulated data in the same way as from the empirical data using *D. procumbens* as a template (i.e., the same number of loci with n SNPs per locus) and two different sequence lengths: 380bp and 147bp. We found that the number of demographic events (“sum(indicators.alltrees)”) was greater than one for all declines simulated at 90% severity, indicating that population size was not constant over time (Table S3). As the severity of the decline decreased, we were unable to reject constant size more often. Constant sizes were consistently rejected at all severities for 147bp simulations, even at smaller simulated changes in population size. This suggests that the “sum(indicators.alltrees)” is not as informative when using shorter sequences. The EBSP-estimated rates of decline were as expected given the original simulated percentage (Fig. 3; Fig. S1). For example, trends estimated using EBSP for 90% contraction consistently exhibited the steepest decline. Estimates of the extent of decline were correlated with simulated proportions at 380bp (Fig. 3; linear regression, p = 0.034, R^2^ = 0.90), but not correlated at 147bp (Fig. S1; p = 0.266, R^2^ = 0.31). We observed a reduction in population size close to the very recent past (<0.005 substitutions/site) regardless of simulated decline severity. In part, this is likely to be due to a lack of signal caused by low numbers of coalescent events near the present. However, this was not observed in the empirical EBSP results for *D. procumbens* (Fig. 2a) and may be an artefact of simulating constant population size (rather than growth) followed by a very recent shift in *N*_*e*_. Nevertheless, these results show that our approach has the power to detect population decline in the Anthropocene and identify the severity of the bottleneck, particularly when using longer sequence lengths.

**Figure 3.**
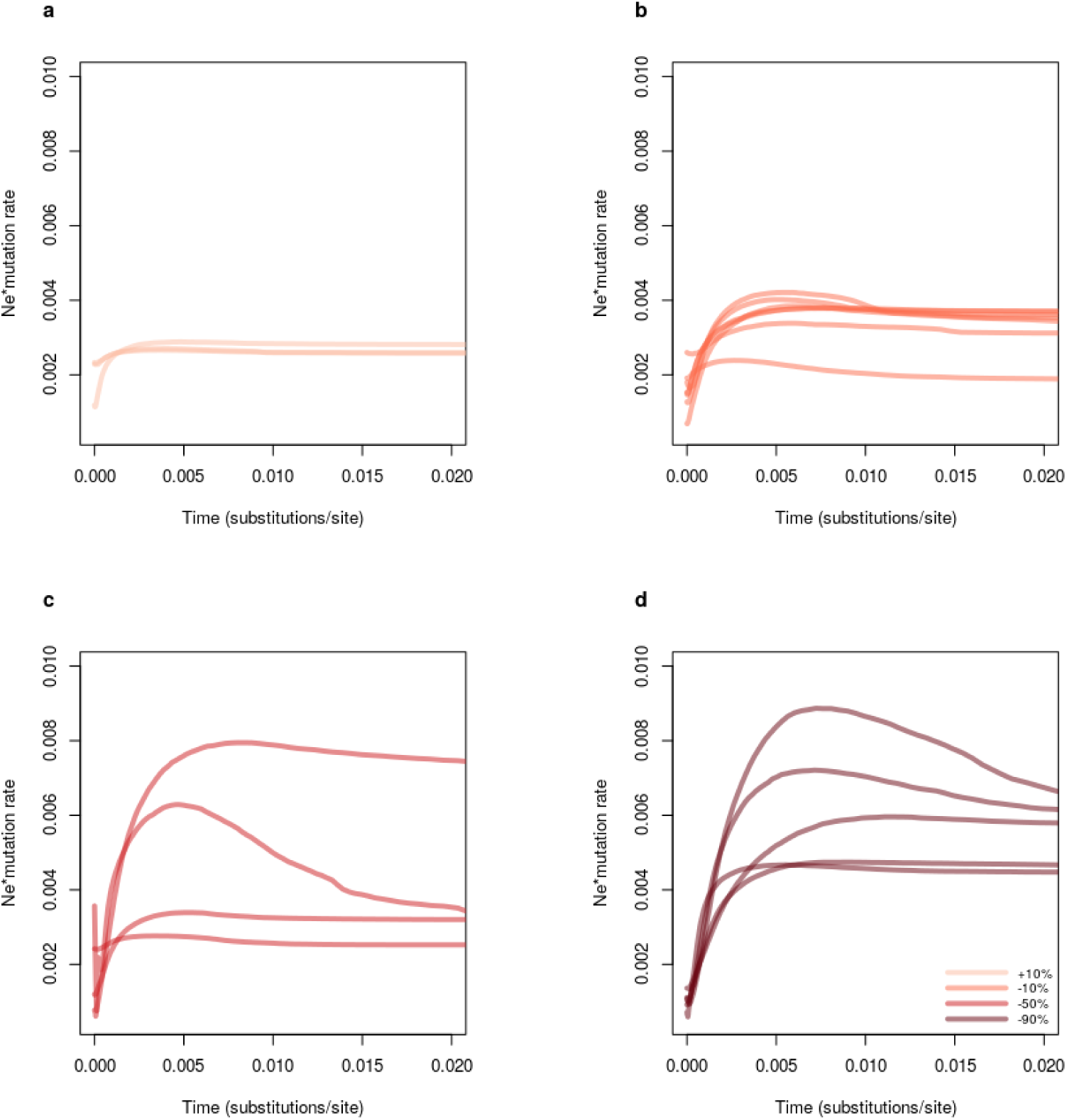
**S**imulated effective population size through time for one expansion (a) +10% and three different levels of population decline (b) -10% (c)-50% (d) -90% at 2500 years before present. We performed simulations using our run for *D. procumbens* as a template, keeping the same number of loci per SNP category as in the empirical run. We performed two sets of simulations: using two different sequences length of 380bp (shown here) and 147bp (shown in Fig. S1). Simulated data was then used in the same pipeline as previously used for empirical data. The x-axis shows time measured in substitutions per site (present on the left side of the graph) and the y-axis shows *N*_*e*_ scaled by mutation rate. Shades of red deepen according to increasing severity of decline.

To investigate the prevalence of these patterns in less well-studied Madagascan species, we repeated the above analyses for the remaining six species in our dataset (Fig. 4) – only one of which has known census population sizes. Among this group, just three species have been red list assessed and are categorised as of least concern: *Psiadia altisimma* and *Dichaetanthera cordifolia* have stable populations (*Psiadia altisimma* is a slash and burn successor), while *Dilobeia thouarsii* is decreasing due to agriculture, harvesting for timber and traditional medicine (Lemmens *et al*. 2012; Letsara 2018). *Chysophyllum boivinianum* is also exploited for timber and malarial medicine, and is suspected to be declining rapidly (Lemmens 2007). Little is known about the extent of exploitation and biology of the other two species in our sample; *Adenia perrieri* and an undescribed species of *Psychotria*. In these six species, *N*_*e*_ ranged from 7,253 in *Psychotria sp*. to 25,702 in *D. cordifolia* (Fig. 4). Although census sizes are not available for these populations, it seems unlikely that any of these species would have always been rare, indeed *P. altisimma* is relatively common (Danthu *et al*. 2008). However, the EBSP analyses are more revealing. Two species showed no evidence of recent decline; *A. perrieri* and, as expected, *D. cordifolia. Psiadia altissima* showed some evidence of decline (↓43%) but we could not reject constant size for this species. The remaining three species showed clear decreases in population size towards the present, which is consistent with the suspected exploitation of *C. boivinianum* (↓83%) and *D. thouarsii* (↓82%). Madagascar is a centre on endemism for *Psychotria* (↓68%), which are often small understory trees and shrubs with limited ranges so these species may be particularly susceptible to deforestation. We compared percentage declines in our empirical data to our simulations (Table S4) and found that empirical declines fell within the range of simulated declines (10%-90% decline) in all cases, but as our simulations have shown there is a greater degree of uncertainty for the analyses where short sequences were used (e.g. *P. altissima* and C. *boivinianum*).

**Figure 4.**
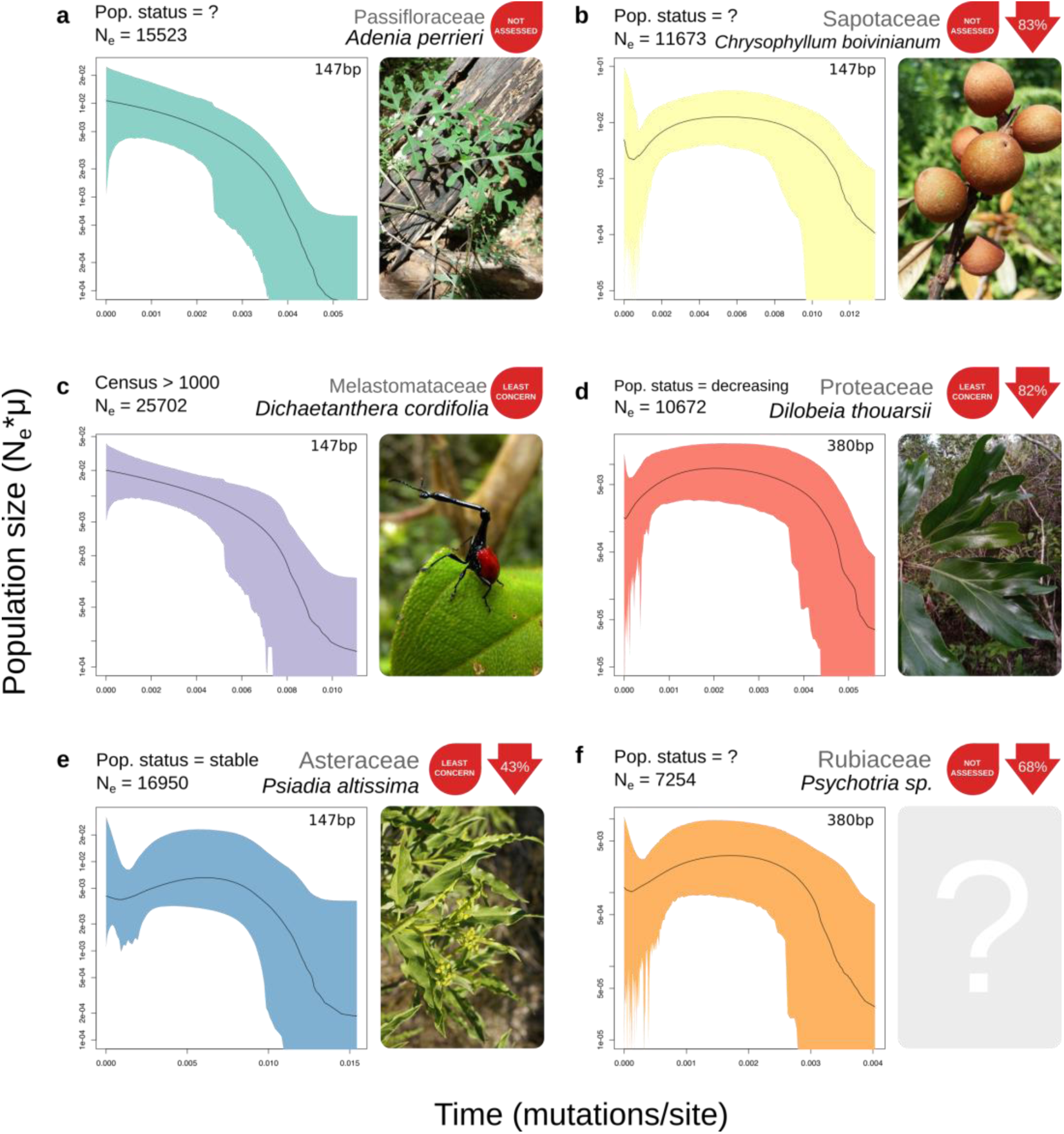
Effective population size through time for six species from six different plant families. The present day is on the left side of each plot. Black lines represent the median population size and shaded polygons show the 95% highest posterior density (HPD). To the right of each plot is a picture of the study species and the current IUCN red list assessment, where available. Values in red arrows indicate the percentage decline from maximum population size to the minimum population size inferred after the maximum. As described in in figure 1, rapid increases towards the present in *C. boivinianum* and *Psychotria sp*. are due to lack of signal. Photos of *A. perrieri, C. boivinianum* and *D. thouarsii* by FR, Solofo Eric Rakotoarisoa and Rahaingoson Fabien respectively and shared under the CC BY-NC 4.0 license (https://creativecommons.org/licenses/by-nc/4.0/). Photo of *D. cordifolia* by Frank Vassen and shared under the CC BY 2.0 license (https://creativecommons.org/licenses/by/2.0/deed.en). Photo of *P. altissima* by B.navez shared under the CC BY-SA 3.0 license (https://creativecommons.org/licenses/by-sa/3.0/). Small modifications have been made to each photo.

Given that demographic decline in the EBSP was only detected in the most severe cases in the palms, the decline observed in these species suggests that they may be at more severe risk of extinction than current knowledge suggests. We were able to repeat EBSP analyses with alternate marker sets in six species (Fig. S2) and found results that were consistent with initial demographic trends (Fig. 2, 4), suggesting that our results are robust to potential marker choice bias. It is important to note, however, that coalescent estimation of population sizes calculated in this way can only determine demographic patterns for the population that the sampled individuals represent, which may not reflect the species as a whole. As the majority of species were collected in the fragmented forests of Andasibe (Fig. 1), it is possible that these declines are only pertinent to the Andasibe populations. This region experienced very rapid deforestation that has recently been limited by community-based conservation efforts (Dolch *et al*. 2015) and our results highlight the high levels of population decline that have happened even in currently well-managed areas.

Our ddRADseq approach is a relatively cheap and accessible way to assess past demographic changes with a handful of specimens rapidly. It holds great potential to screen species at a much higher rate than is possible with observation-based red listing assessments. A single protocol was successful for species from seven different plant families, highlighting the universality of our approach. Demographic trends can be inferred using just a few individuals with large numbers of markers (Heled & Drummond 2008), where previous approaches with small numbers of markers (e.g., microsatellites or chloroplast DNA) fall short. This makes our approach ideal for studying rare and difficult to access species, many of which may be the most threatened and in need of assessment. Effective population sizes estimated using coalescent methods are not proxies for the variance effective size often used in conservation efforts (Lauterbur 2019), but changes in *N*_*e*_ are informative regarding recent demographic changes, as is shown here. Further work is required to establish the nature of the relationship between the population declines observed in EBSP analyses and the extinction risk of these species, and to determine what role these types of analyses can play in red listing efforts. Our results suggest that rapid decline has affected 70% of the species we assessed and it seems likely that many of the micro-endemic species in Madagascar have not always been so rare. Instead, we propose that the radical environmental changes that have taken place in Madagascar since the arrival of humans have driven rapid population decline in Madagascan plants.

## Materials and Methods

### Sample Collection

43 individuals across 10 species and seven families were sampled (Table S1). Leaf tissue was collected from wild individuals and desiccated using silica gel to preserve DNA. Approximately 20mg of dried tissue was used for DNA extractions. The majority of samples were collected in Andasibe, Madagascar and palm species were collected across the island (Fig. 1). Location data are detailed in Table S1.

### Library preparation

Genomic DNA was extracted using a hexadecyltrimethylammonium bromide (CTAB) mini-extraction protocol (Doyle & Doyle 1987), purified using spin columns from the Qiagen DNeasy Plant Mini Kit and then eluted in 60μl water. Double-digest Restriction-site Associated DNA (ddRAD) libraries were constructed following the protocol of Peterson et al. (Peterson *et al*. 2012). Briefly, 1 μg of high molecular weight DNA was digested using two rare-cutting enzymes; EcoRI-HF (NEB) and SphI-HF (NEB). Barcoded flex adapters (Peterson *et al*. 2012) were ligated to 400 ng digested DNA for multiplexing and samples were pooled. Size selection was carried out with a Pippin Prep (Sage Biosciences) with a tight window of 468 to 546bp. Eight PCR reactions per library were conducted to minimize the effect of PCR bias.

### Sequencing and genotyping

Libraries were sequenced on three Illumina MiSeq runs, using 2×75bpb (V3), 2×250bp (V2) and 2×300bp (V3) kits respectively. Read pairs were concatenated to produce a single sequence of either 147bp or 380bp after barcode removal and trimming. Reads were demultiplexed using *process_radtags* in the software Stacks (v1.40) (Catchen *et al*. 2013). Reads were aligned and SNPs detected using *ustacks*. A separate catalogue was built for each species (*cstacks*), to which loci were matched from each individual of a species (*sstacks*). *Populations* was used to filter and output data by only retaining those loci that were present in all individuals within a species.

### Inference of *N*_*e*_ and demographic history

For each species, the fasta format output of from the STACKS pipeline containing all aligned loci, was used to estimate effective population size. We first estimated the population scaled mutation rate (*θ* = 4*N*_*e*_*µ*) using a Bayesian method, which permits for rate variation between loci, implemented in the R package *thetamateR*. The estimated *θ* value for each species was then converted to *N*_*e*_ assuming an average silent site mutation rate in angiosperms of 5.35 × 10 ^−9^ sites/year (De La Torre *et al*. 2017) and a generation time of 10 years.

To infer changes if population size through time, we used the EBSP (Heled & Drummond 2008) approach in BEAST v2.4.2 (Bouckaert *et al*. 2014), following the methodology outlined in Trucchi et al. (Trucchi *et al*. 2014) for each species separately. A custom R script (Trucchi *et al*. 2014) was used to select only those loci with high numbers of SNPs (3 or more per locus) to maximise the amount of information per marker to increase the accuracy and reliability of our inference. From these 50 loci were selected randomly to use in the EBSP analysis. One haplotype was chosen at random for each individual at heterozygous loci to minimize bias in haplotype frequencies and reduce bias caused by undiscovered heterozygote samples (Trucchi *et al*. 2014). A separate substitution model was assigned for each SNP class with kappa linked between models, trees were unlinked across loci and a single strict clock was assigned to all loci. Operators were modified to increase mixing efficiency (Trucchi *et al*. 2014). Each MCMC ran for 500,000,000 generations, sampling every 50,000 and the level of stationarity (*R. robustior* was run for twice this length). Stationarity was verified using Tracer v1.6 (Drummond & Rambaut 2007). Multiple replicate runs were conducted to ensure the same stationarity point was reached and runs combined to increase ESS values to suitable levels (ESS > 100 for all important parameters). EBSPanalyser in BEAST was applied to perform a linear reconstruction with data from combined runs. If an additional, unused set of 50 highly-variable loci was available analyses were repeated and results compared across datasets.

### Simulated data

We used fastsimcoal2 (v2.6) (Excoffier et al. 2013) to simulate sequence data for a genetic bottleneck 250 generations ago (equivalent to 2500 years). We simulated four strengths of bottleneck; a 90%, 50%, 10% reduction and a 10% increase in effective population size with ancestral population set to our thetamateR *N*_*e*_ estimates *D. procumbens*. We generated 40,000 DNA sequences matching the length of the corresponding empirical RAD tags (147bp, 380bp) and selected loci with the same distribution of SNPs in the empirical *D. procumbens* dataset (i.e., each SNP category contained the same number of tags). We then used these loci as input for our pipeline (described above). We repeated simulations for each decline severity at least three times to ensure consistency in demographic patterns across different simulations.

## Acknowledgments

We thank the Howard Lloyd Davies legacy and NERC for funding. Rhian J. Smith, Simon Creer, Andrew Foote and Kathy Willis for support and comments.

## Author Contributions

ASTP conceived and designed the study with input from all authors. AJH generated and analysed the data with contributions from ASTP. SC coordinated sample collection by RR and FR. WLE and WJB provided samples. AJH and ASTP wrote the manuscript with contributions from all authors.

