## Supplementary material for "The Demographic History of Micro-endemics: Have Rare Species Always Been Rare?": SI

**Figure S1** Simulated extended Bayesian skyline plots (EBSPs).
**Figure S2** Second runs from Extended Bayesian Skyline Plot

**Table S1** Details of specimens used in this study
**Table S2** Sequencing results and summary statistics.
**Table S3** Number of demographic events in our simulations
**Table S4** Percentage declines in empirical and simulated EBSPs

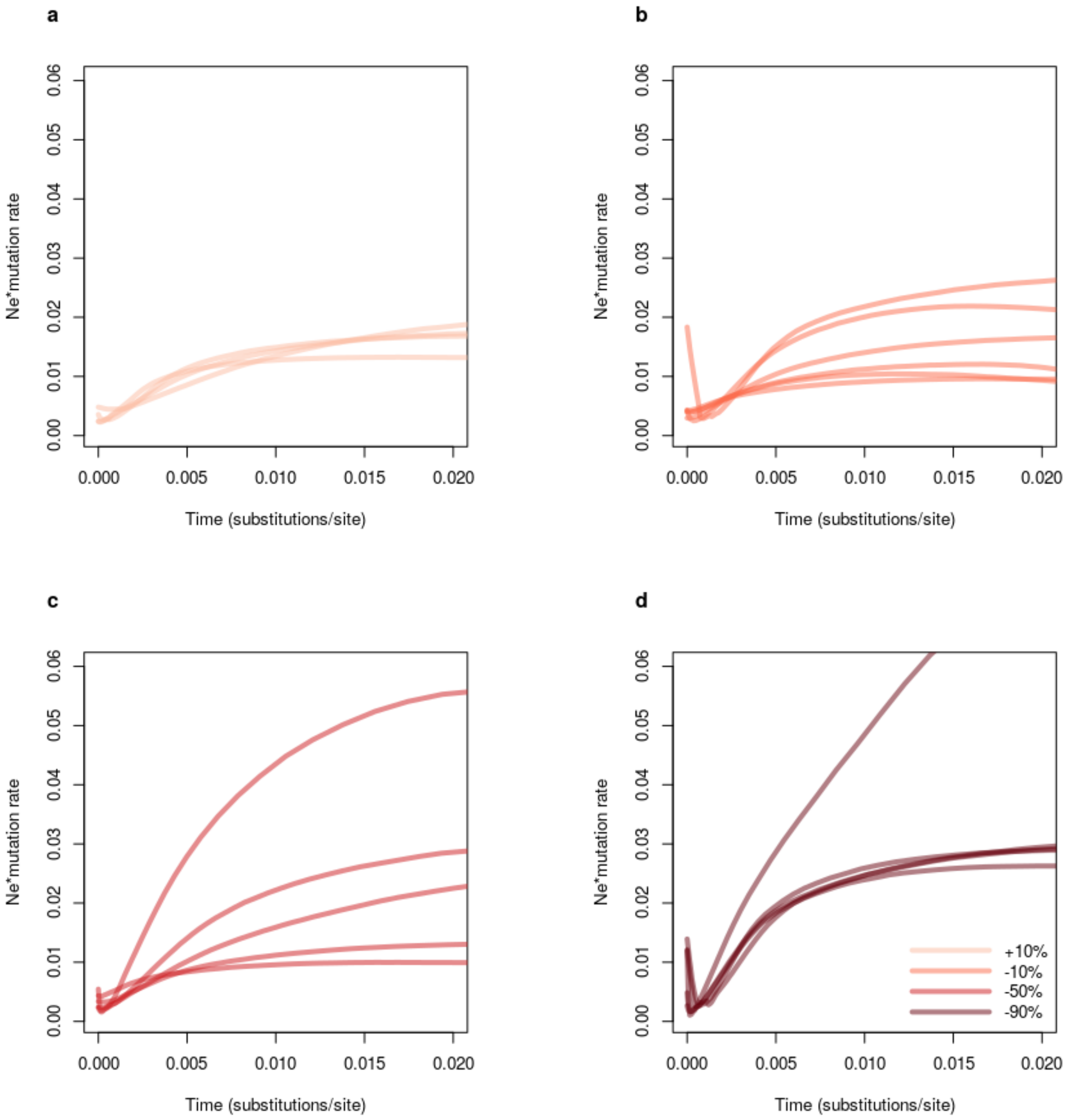


**Figure S1** Simulated effective population size through time for one expansion (a) +10% and three different levels of population decline (b) -10% (c)-50% (d) -90% at years before present. We performed simulations using our run for *D. procumbens* as a template, keeping the same number of loci per SNP category as in the empirical run. We performed two sets of simulations: using two different sequences length of 380bp (shown in Fig. 3) and 147bp (shown here). Simulated data was then used in the same pipeline as previously used for empirical data. The x-axis shows time measured in substitutions per site (present on the left side of the graph) and the y-axis shows Ne scaled by mutation rate. Shades of red deepen according to increasing severity of decline.

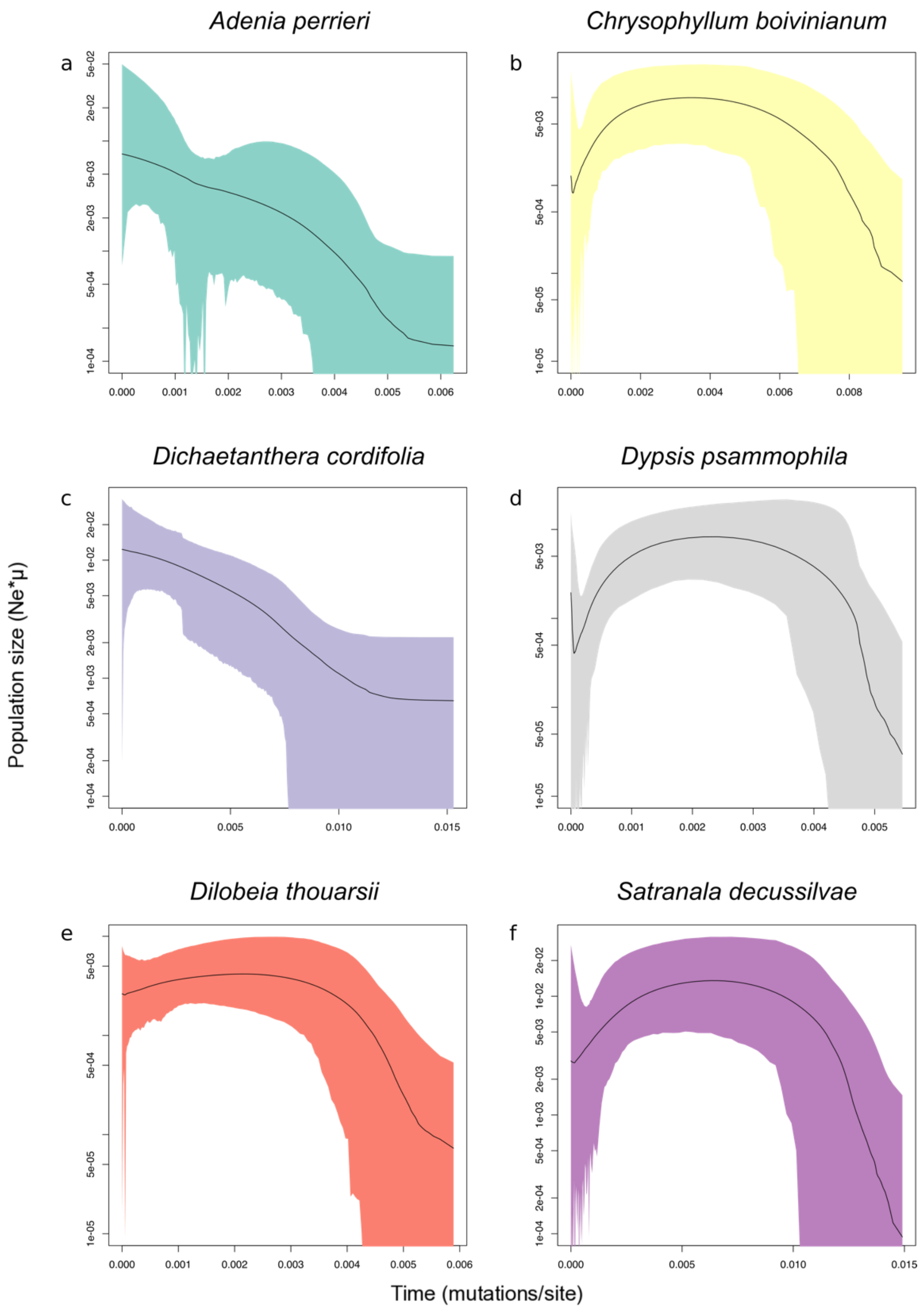


**Figure S2** Second runs from Extended Bayesian Skyline Plot analyses showing $N_e$ through time. Each run was performed with an entirely different marker set compared to those in figures 2 and 3. Black lines represent median values, shaded polygons represent 95% Highest Posterior Density (HPD) values inferred for population size.

**Table S1** Details of specimens used in this study. Taxonomic information, code and location information are shown where available.

| Family | Genus | Species | Code | Longitude | Latitude |
|---|---|---|---|---|---|
| Passifloraceae | *Adenia* | *perrieri* | Ap1 | -18.804722 | 48.428333 |
| Passifloraceae | *Adenia* | *perrieri* | Ap2 | -18.804722 | 48.428333 |
| Passifloraceae | *Adenia* | *perrieri* | Ap3 | -18.804722 | 48.428333 |
| Passifloraceae | *Adenia* | *perrieri* | Ap4 | -18.804722 | 48.428333 |
| Sapotaceae | *Chrysophyllum* | *boivinianum* | Cb1 | -18.804722 | 48.428333 |
| Sapotaceae | *Chrysophyllum* | *boivinianum* | Cb2 | -18.804722 | 48.428333 |
| Sapotaceae | *Chrysophyllum* | *boivinianum* | Cb3 | -18.804722 | 48.428333 |
| Sapotaceae | *Chrysophyllum* | *boivinianum* | Cb4 | -18.804722 | 48.428333 |
| Sapotaceae | *Chrysophyllum* | *boivinianum* | Cb5 | -18.804722 | 48.428333 |
| Melastomataceae | *Dichaetanthera* | *cordifolia* | Dc1 | -18.804722 | 48.428333 |
| Melastomataceae | *Dichaetanthera* | *cordifolia* | Dc2 | -18.804722 | 48.428333 |
| Melastomataceae | *Dichaetanthera* | *cordifolia* | Dc3 | -18.804722 | 48.428333 |
| Melastomataceae | *Dichaetanthera* | *cordifolia* | Dc4 | -18.804722 | 48.428333 |
| Melastomataceae | *Dichaetanthera* | *cordifolia* | Dc5 | -18.804722 | 48.428333 |
| Proteaceae | *Dilobeia* | *thouarsii* | Dt1 | -18.804722 | 48.428333 |
| Proteaceae | *Dilobeia* | *thouarsii* | Dt2 | -18.804722 | 48.428333 |
| Proteaceae | *Dilobeia* | *thouarsii* | Dt3 | -18.804722 | 48.428333 |
| Proteaceae | *Dilobeia* | *thouarsii* | Dt4 | -18.804722 | 48.428333 |
| Proteaceae | *Dilobeia* | *thouarsii* | Dt5 | -18.804722 | 48.428333 |
| Arecaceae | *Dypsis* | *cf. psammophila* | WLE128b (Dpsa5) | -15.746667 | 50.182778 |
| Arecaceae | *Dypsis* | *cf. psammophila* | WLE137 (Dpsa1) | -15.728056 | 50.213889 |
| Arecaceae | *Dypsis* | *cf. psammophila* | WLE138 (Dpsa2) | -15.728056 | 50.213889 |
| Arecaceae | *Dypsis* | *cf. psammophila* | WLE139 (Dpsa3) | -15.728056 | 50.213889 |
| Arecaceae | *Dypsis* | *procumbens* | 356 (Dpro4) | -23.536389 | 47.081667 |
| Arecaceae | *Dypsis* | *procumbens* | 7755 (Dpro3) | -18.879167 | 48.47 |
| Arecaceae | *Dypsis* | *procumbens* | WLE72 (Dpro1) | -21.330833 | 47.703056 |
| Arecaceae | *Dypsis* | *procumbens* | WLE77 (Dpro 2) | -21.406944 | 47.945833 |
| Asteraceae | *Psiadia* | *altissima* | Pa1 | -18.804722 | 48.428333 |
| Asteraceae | *Psiadia* | *altissima* | Pa2 | -18.804722 | 48.428333 |
| Asteraceae | *Psiadia* | *altissima* | Pa3 | -18.804722 | 48.428333 |
| Asteraceae | *Psiadia* | *altissima* | Pa4 | -18.804722 | 48.428333 |
| Asteraceae | *Psiadia* | *altissima* | Pa5 | -18.804722 | 48.428333 |
| Rubiaceae | *Psychotria* | *sp.* | Psy1 | -18.804722 | 48.428333 |
| Rubiaceae | *Psychotria* | *sp.* | Psy2 | -18.804722 | 48.428333 |
| Rubiaceae | *Psychotria* | *sp.* | Psy3 | -18.804722 | 48.428333 |
| Rubiaceae | *Psychotria* | *sp.* | Psy4 | -18.804722 | 48.428333 |
| Rubiaceae | *Psychotria* | *sp.* | Psy5 | -18.804722 | 48.428333 |
| Arecaceae | *Ravenea* | *robustior* | WLE116(Rr1) | -14.438889 | 49.752778 |
| Arecaceae | *Ravenea* | *robustior* | WLE361 (Rr2) | -21.836667 | 47.903889 |
| Arecaceae | *Satranala* | *decussilvae* | AA71a (Sd1) | -15.727222 | 50.213333 |
| Arecaceae | *Satranala* | *decussilvae* | AA71j (Sd2) | -15.727222 | 50.213333 |
| Arecaceae | *Satranala* | *decussilvae* | MAN1 (Sd3) | -16.6785134 | 49.739068 |
| Arecaceae | *Satranala* | *decussilvae* | MAN2 (Sd4) | -16.6785134 | 49.739068 |

**Table S2** Sequencing results and summary statistics. Number of reads is the total reads generated from all individuals in a species calculated after cleaning. Number of loci represents the number of loci found in at least one individual. Shared loci details the number of shared markers between all individual. Measures of nucleotide diversity (Pi), heterozygosity and the inbreeding coefficient ($F_{is}$) were taken from those loci shared between all individuals in a species including variant and invariant sites. Mean events shows the mean number of events inferred in the EBSP analyses or the sum(indicator.alltrees) parameter followed by the 95% highest posterior density minimum and maximum. If multiple locus sets were used in EBSP analyses, the statistics from the second run are also shown.

| Species | n | Read length | No. reads | Loci | Shared loci | π | Ho | He | $F_{IS}$ | Mean events [95% HPD] |
|---|---|---|---|---|---|---|---|---|---|---|
| *Adenia perrieri* | 4 | 147 | 5531474 | 26921 | 2476 | 0.3358 | 0.2969 | 0.2938 | 0.0828 | 1.402 [1,3], 1.5726 [1,4] |
| *Chrysophyllum boivinianum* | 5 | 147 | 9714612 | 26736 | 1733 | 0.3361 | 0.2976 | 0.3025 | 0.0921 | 2.998 [2,4], 2.798 [2,4] |
| *Dichaetanthera cordifolia* | 5 | 147 | 12836869 | 48565 | 1883 | 0.309 | 0.2644 | 0.2781 | 0.1076 | 1.444 [1,3], 1.534 [1,3] |
| *Dilobeia thouarsii* | 5 | 380 | 10477774 | 24566 | 3879 | 0.3205 | 0.3196 | 0.2884 | 0.004 | 2.544[2,4], 2.274 [1,4] |
| *Dypsis procumbens* | 4 | 380 | 6983552 | 53520 | 316 | 0.3265 | 0.192 | 0.2856 | 0.296 | 1.182 [1,2] |
| *Dypsis rabepierrei* | 4 | 380 | 9287523 | 24545 | 7372 | 0.4161 | 0.3446 | 0.3641 | 0.1476 | 2.883 [2,4], 2.986 [2,4] |
| *Psiadia altissima* | 5 | 147 | 8689114 | 42583 | 1366 | 0.3116 | 0.2893 | 0.2805 | 0.0565 | 2.492 [0,4] |
| *Psychotria sp.* | 5 | 380 | 7575095 | 48620 | 562 | 0.3436 | 0.2283 | 0.3092 | 0.3138 | 2.696 [1,4] |
| *Ravenea robustior* | 2 | 598 | 4364546 | 51714 | 760 | 0.5293 | 0.4541 | 0.397 | 0.1128 | 1.415 .047 [1,1] |
| *Satranala decussilvae* | 4 | 147 | 8386955 | 29088 | 6100 | 0.3674 | 0.289 | 0.3215 | 0.1505 | 2.628 [2,4], 2.7 [2,4] |

**Table S3** Number of demographic events in our simulations. We examined the mean and 95% highest posterior densities (HPDs) of the 'sum(indicators.alltrees)' parameter from our EBSP runs. If the 95% HPD minimum does not overlap with zero then we can reject the hypothesis of constant population size over time.

| % decline | sequence length (bp) | Mean | lower | upper |
|---|---|---|---|---|
| 0.5 | 147 | 1.758720284 | 1 | 3 |
| 0.5 | 147 | 1.7700511 | 1 | 3 |
| 0.5 | 147 | 1.415018885 | 0 | 3 |
| 0.5 | 147 | 1.895134415 | 1 | 3 |
| 0.5 | 147 | 2.658192969 | 1 | 4 |
| 0.5 | 380 | 2.637882761 | 1 | 4 |
| 0.5 | 380 | 1.580537658 | 0 | 3 |
| 0.5 | 380 | 0.845145523 | 0 | 3 |
| 0.5 | 380 | 1.874472339 | 1 | 3 |
| 0.1 | 147 | 1.395356587 | 0 | 3 |
| 0.1 | 147 | 1.908575872 | 0 | 4 |
| 0.1 | 380 | 1.442679405 | 0 | 3 |
| 0.1 | 380 | 1.393245945 | 0 | 3 |
| 0.1 | 147 | 1.710842035 | 1 | 3 |
| 0.1 | 147 | 2.10330652 | 0 | 4 |
| 0.1 | 147 | 2.28682515 | 1 | 4 |
| 0.1 | 147 | 2.461779476 | 1 | 4 |
| 0.1 | 380 | 1.811153077 | 0 | 3 |
| 0.1 | 380 | 1.14419018 | 0 | 3 |
| 0.1 | 380 | 1.05376583 | 0 | 3 |
| 0.1 | 380 | 1.771384137 | 0 | 4 |
| 0.1 | 380 | 1.695289936 | 1 | 3 |
| 0.9 | 147 | 1.986111728 | 1 | 3 |
| 0.9 | 147 | 1.9508998 | 1 | 3 |
| 0.9 | 147 | 2.162741613 | 1 | 4 |
| 0.9 | 147 | 2.836369696 | 1 | 4 |
| 0.9 | 147 | 2.315707621 | 1 | 4 |
| 0.9 | 380 | 1.648189291 | 1 | 3 |
| 0.9 | 380 | 1.579093535 | 1 | 3 |
| 0.9 | 380 | 2.162659437 | 1 | 4 |
| 0.9 | 380 | 2.008442568 | 1 | 4 |
| 0.9 | 380 | 1.809264608 | 1 | 3 |
| 1.1 | 147 | 1.835147745 | 1 | 3 |
| 1.1 | 147 | 1.94345701 | 1 | 3 |
| 1.1 | 147 | 1.920559086 | 1 | 4 |
| 1.1 | 147 | 1.711508554 | 1 | 3 |
| 1.1 | 380 | 0.774161297 | 0 | 3 |
| 1.1 | 380 | 1.398911353 | 0 | 3 |
| 1.1 | 380 | 0.830926461 | 0 | 3 |

**Table S4** Percentage declines in empirical and simulated extended Bayesian skyline plots (EBSPs). We identified the maximum effective population size ($N_e$) and the minimum $N_e$ after the maximum had been reached and used these to calculate percentage decline (see Fig. 1). Declines for simulations were calculated after removing the most recent 12.5% towards the present to avoid bias caused by sudden decreases/increases in $N_e$ due to missing data that was common to almost all simulation-based EBSPs. We then averaged the decline across runs for each read length/simulated decline. Zeros indicate that a decline was not modelled in that species' history.

| species | read length | Simulated decline | % decline |
|---|---|---|---|
| simulated | 380 | 1.1 | 0.2669692 |
| simulated | 380 | 0.1 | 0.5107119 |
| simulated | 380 | 0.5 | 0.598272 |
| simulated | 380 | 0.9 | 0.8324882 |
| simulated | 147 | 1.1 | 0.825631 |
| simulated | 147 | 0.1 | 0.7315994 |
| simulated | 147 | 0.5 | 0.8185062 |
| simulated | 147 | 0.9 | 0.935723 |
| *Adenia perrieri* | 147 | | 0 |
| *Chrysophyllum boivinianum* | 147 | | 0.8271917 |
| *Dichaetanthera cordifolia* | 147 | | 0 |
| *Dilobeia thouarsii* | 380 | | 0.8214475 |
| *Dypsis procumbens* | 380 | | 0 |
| *Dypsis rabepierrei* | 380 | | 0.8966892 |
| *Psiadia altissima* | 147 | | 0.4310314 |
| *Psychotria sp.* | 380 | | 0.6780376 |
| *Ravenea robustior* | 598 | | 0 |
| *Satranala decussilvae* | 147 | | 0.7787335 |
